# Disentanglement of batch effects and biological signals across conditions in the single-cell transcriptome

**DOI:** 10.1101/2025.04.10.648296

**Authors:** Shunta Sakaguchi, Masato Tsutsumi, Kentaro Nishi, Honda Naoki

## Abstract

Batch effects, which arise due to technical variations across different experimental factors, pose a significant challenge for single-cell transcriptomic analysis. Although various batch correction methods have been developed to mitigate these effects, they often indiscriminately mix data from different batches, leading to the removal of biologically meaningful signals. This limitation hinders comparative analyses across multiple conditions, an essential aspect of scientific research. Recent approaches attempt to address this issue by mapping data to separate spaces, but they prevent direct comparisons between conditions. Here, we propose Kanade, a batch correction method based on a variational autoencoder. Kanade explicitly disentangles batch effects from biological signals by specializing latent variables for different types of information. Using both simulated and real datasets, we demonstrate that Kanade selectively correct batch effects while preserving essential biological features, enabling more accurate comparative analyses in single-cell transcriptomics.

## Introduction

Single-cell omics is an essential technology for investigating the mechanisms of various biological processes including cell differentiation [1] and diseases [2, 3]. Recently, a large number of single-cell transcriptomic datasets have been obtained [4, 5], and most of these datasets consist of multiple batches. Among the batches, unwanted variations (batch effects) arise due to differences in technical platform, experimenter, or other factors and often confound biological signals [6]. This is an obstacle to analyzing the single-cell transcriptome dataset.

To overcome the problem, many correction methods for the batch effect have been developed. For example, Seurat [7, 8] has a data integration tool, and Harmony [9] shows high performance in correcting batch effects [10]. In addition, variational autoencoder (VAE)-based methods including scVI [11] are often used. However, these methods attempt to mix data from different batches as much as possible and eliminate biological signals between different conditions (e.g. health and disease) which should be preserved (Fig. 1A, center-right). Since comparisons between multiple conditions are fundamental to scientific research, this issue severely limits single-cell transcriptome analyses. Although methods for comparison conditions such as multiGroupVI [12] have been proposed in recent years, they map data to separate latent-variable spaces for each condition, making direct comparisons impossible.

**Fig. 1:**
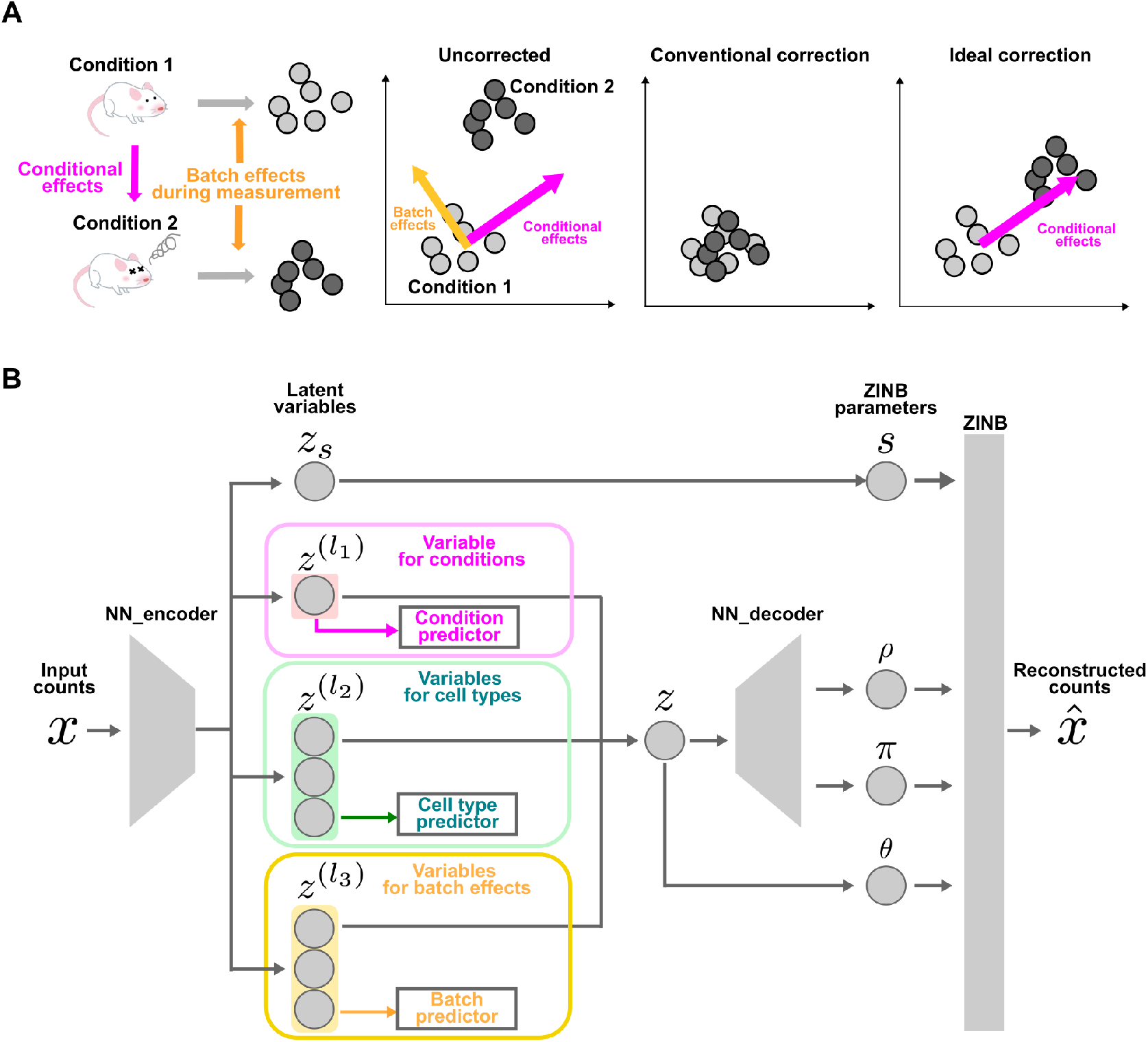
Overview of Kanade. (A) A schematic illustration of the confounding of conditional effects and batch effects. Each sample obtained from different conditions is confounded by batch effects (left). The observed difference reflects the combined effects of both factors (center-left). Conventional correction methods remove both the conditional effect and the batch effect (center-right). Ideally, only the batch effect should be selectively removed while preserving the conditional effect (right). Illustrations of mice in the figure were adapted from materials by Irasutoya (B) Schematic diagram of Kanade model structure.

A common issue with existing methods is that they do not explicitly separate confounding batch effects from biological signals. One solution for the issue is to constrain sets of latent variables to assign specific roles to them. This assignment has been used in the extraction of morphological information from images. For instance, Morpho-VAE [13] controls the latent space of the VAE to preserve morphological information for species classification by simultaneously training predictors that infer species from the latent variables.

Here, we propose a batch correction method called Key Approach for Noise Adjustment and DisEntanglement (Kanade; Fig. 1B). The most distinctive feature of this tool is that it is designed to disentangle batch effects in transcriptomic data from biological signals between different conditions. Kanade is based on the variational autoencoder and specializes parts of latent variables in capturing specific types of information using predictors. This makes it possible to separate batch effects from biological signals and to correct batch effects by replacing only the latent variables for batch effects with zeros or other uniform values. First, we demonstrated that Kanade was able to selectively correct batch effects and conserve conditional features using simulated datasets. Furthermore, Kanade was applied to the scRNA-seq atlas of pancreatic cells and corrected batch effects without losing diabetes-related features.

## Result

### The Kanade model

Kanade is a VAE-based model (Fig. 1B). In the Kanade model, the expression *x*_*i,g*_ of gene *g* in cell *i* is assumed to be sampled from a zero-inflated negative binomial distribution (ZINB). ZINB is parameterized by the scale factor *s*, the normalized mean *ρ*, the dispersion parameter *θ*, and the zero-inflated rate *π*. Kanade uses a neural network to encode *x*_*i*_, a gene expression vector in cell *i*, into a latent variable vector *z*_*i*_ and a latent variable for scale factor 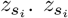 is converted into the scale factor *s*, and *z*_*i*_ is decoded into other ZINB parameters.

To disentangle batch effects from biological signals across different conditions, Kanade employs specific predictors. The latent variable space is partitioned into distinct sets of variables: one set is designated to predict batch labels for individual cells, while several other sets are assigned to predict the respective condition labels. The remaining set is dedicated to predicting cell types. Then, The entire model is trained so that the likelihood of ZINB for the input gene expression vector and predicitivity of each predictor are jointly maximized. This makes each latent variable retain corresponding information to the predictor to which it is assigned.

Specification of latent variables enables the correction of batch effects. First, data are projected into latent variables, and batch effects and signals depending on condition are stored in the corresponding set of latent variables. Then, replacing only latent variables for batch effects by zeros and reconstructing gene expressions result in corrected gene expressions.

### Biological signals and batch effects were disentangled in latent space on the simulated datasets

To evaluate Kanade, we generated a simulated dataset (Methods) and applied Kanade to it. The dataset has discrete wild-type (WT) / Disease conditions (Discrete data, Fig. 2A). The dataset included three cell types. Batch effects were randomly introduced in each batch(Figs. 2A-C).

**Fig. 2:**
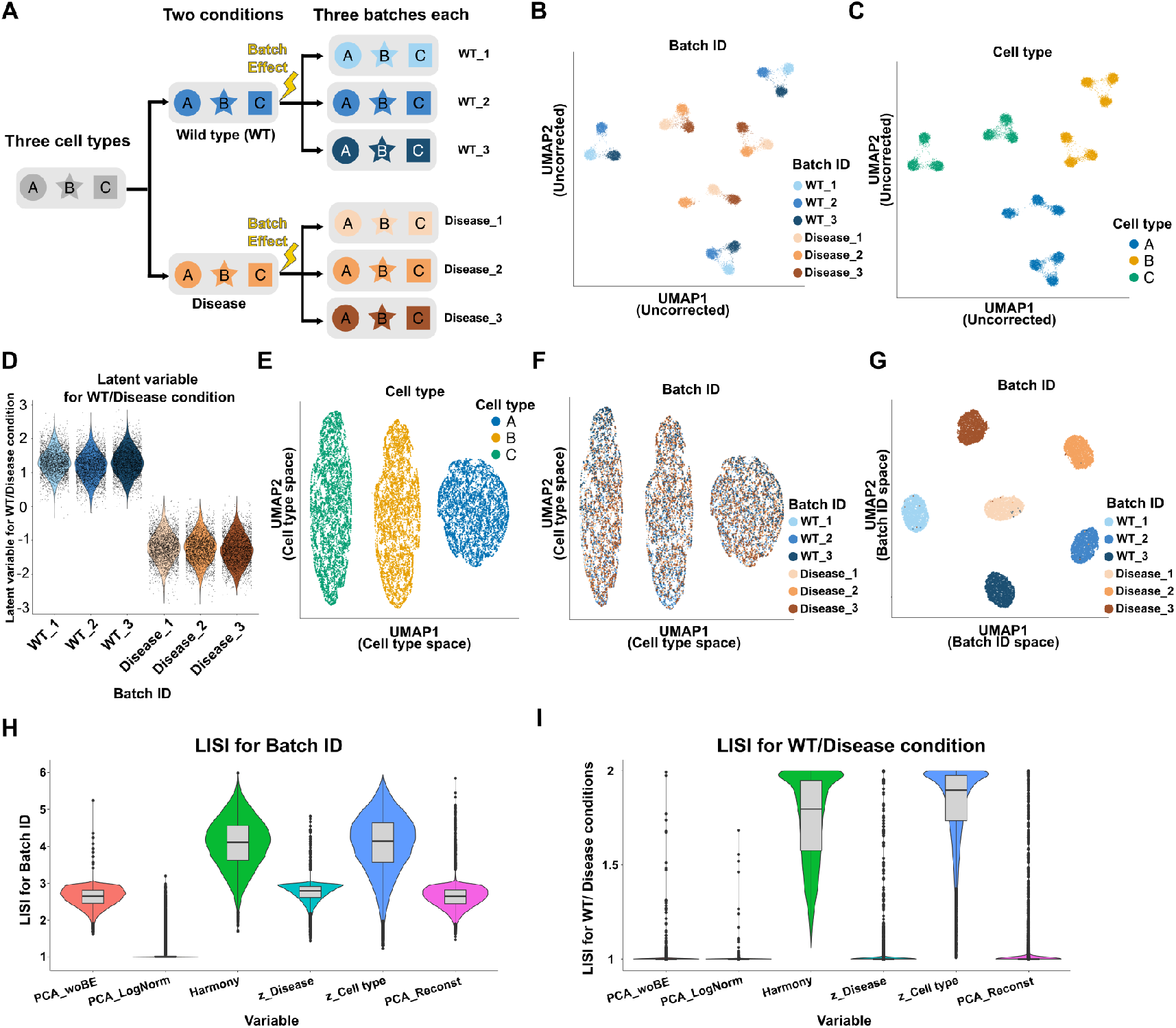
Simulated discrete data in the latent variable spaces of Kanade. (A) The scheme of discrete data.(B and C) UMAP plots of discrete data based on log-normalization. Colors of dots show Batch IDs (B) and cell types (C). (B) The violin plots of the latent variable for WT/Disease condition across independent batches. Each dot represents an individual cell. (E and F) UMAP plots of discrete data using latent variables for cell types. Colors of dots show cell types and batch IDs (F). (G) UMAP plot of discrete data using latent variables for batches. Colors of dots show batch IDs. (H and I) The violin plots and box plots of LISI for Batch ID (H) and WT/Disease conditions (I). Dots represent outliers in box plots.

First, using Discrete data, we evaluated whether Kanade can separate biological signals and batch effects in the latent variable spaces. The latent variables for the WT / Disease condition show clear differences between the two conditions, whereas minimal variation among batches within the same condition (Fig. 2D). Moreover, three cell types were clearly clustered in the latent variable space for cell types, and all batches were mixed in each cluster (Fig. 2D and E). Furthermore, in the latent variable space for batches, all batches were clearly distinguished (Fig. 2F). These results indicate that the three types of information were distinctly disentangled within the latent variable space.

To evaluate these separations quantitatively, we calculated local inverse Simpson’s indices (LISI) [9] for Batch IDs (Fig. 2H) in each latent variable space. Inverse Simpson’s index represents the effective number of mixed components within a population, and LISI extends this concept by computing it at the single-cell level and represent the effective number of mixed components around a cell. In the absence of batch effects, where each condition contains three batches, LISI values approached 3.0 for most of the cells in the principal conponent analyse (PCA) space (PCA woBE). This means that three batches in each condition were mixed around most of cells. When batch effects were introduced, the cells were segregated by batch, resulting in LISI values of approximately 1 for nearly all cells (PCA LogNorm). This means that cells were separated into clusters consisting of single batches. In the latent variable for Disease by Kanade, LISI values were around 3.0, indicating three batches of each condition were almost perfectly mixed (z Disease). LISI in the latent variables for cell types (z cell type) by Kanade showed comparable (median 4.14) to those in Harmony-corrected variables (median 4.11) indicating a comparable level of batch effect correction to that achieved by Harmony, which has high performance of the batch correction [10]. LISIs for WT/ Disease conditions were also calculated (Fig. 2I). LISIs for WT/ Disease in z Disease showed nearly 1 as well as in PCA woBE and PCA LogNorm, indicating a clear separation of two conditions. In contrast, LISIs for WT/ Disease in z cell type almost reached 2.0 as well as Harmony. These results quantitatively demonstrated the ability of Kanade to remove batch effects while preserving condition-specific information in the latent variable space.

To strengthen the validation, we constructed a dataset based on the assumption of continuouse-timedependent gene expression changes and cell sampling at five time points (Continuous data, Figs. S1A-C). When Kanade was applied to the dataset, cells were separated in each latent variable space based on the corresponding signal (Figs. S1D-G). LISIs for Batch ID in latent variables for time point (z TP in Fig. S1H) showed nearly 3, indicating that batches in each time point were mixed, as well as those in latent variables for cell type (z Cell type in Fig. S1H) showed comparable values to those in Harmony-corrected latent variables. LISIs for time points in z TP of most cells were nearly 1, indicating separation depending on time, and those in z Cell type showed comparable values to Harmony (Fig. S1G).

### Batch effects were corrected in reconstructed counts

To test whether Kanade is able to remove batch effects in reconstructed counts and keep biological signals, the latent variables for batch effects in Discrete data were replaced by zeros and counts were reconstructed using these corrected latent variables under the uniform scale factors. First, mean reconstructed counts of genes were highly correlated with the ground truth (Fig. 3A). This suggested that the expression profiles of most genes were accurately recovered. Next, dimensionality reduction was performed on reconstructed counts. Cells were clustered depending on cell types and WT / Disease conditions on the UMAP plot of reconstructed counts without batch-specific segregation within conditions (Figs. 3B and C). LISIs were calculated in PCA space based on the reconstructed counts (PCA reconst in Figs. 2H and I). The LISIs for Batch IDs exhibited a distribution similar to that observed in the absence of batch effects, while the LISIs for WT/Disease conditions were close to 1 in most cases. These results indicated that batches were mixed in the same conditions and differences between conditions were preserved in reconstructed counts.

**Fig. 3:**
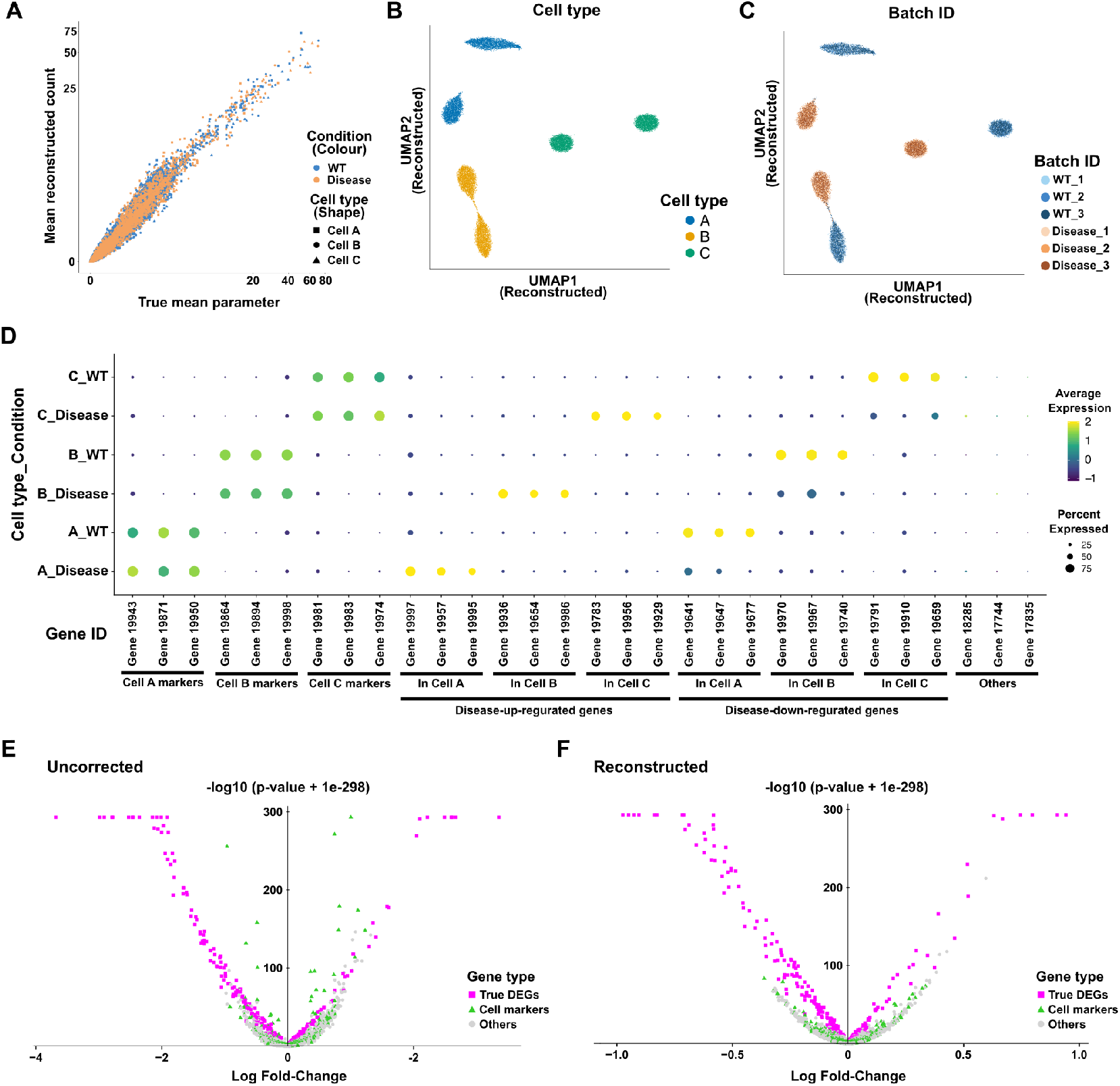
Reconstructed counts of simulated discrete data. (A) Scatter plots of mean gene expressions in each cell type and condition. The X -axis represents the mean parameters of the simulation and the Y-axis mean of reconstructed counts. Both axes are log-scaled. Each dot represents each gene, and the color and shape of the dots represent conditions and cell types respectively. (B and C) UMAP plots of discrete data based on reconstructed counts. The colors of the dots show cell types (B) and batch IDs (C). (D) The dot plot of reconstructed gene expressions in each cell type and condition. (E and F) Volcano plots of differentially expressed gene analysis in cell type A between WT and Disease conditions results based on uncorrected log-normalization (E) and reconstructed counts by Kanade (F). Each dot means a gene and X-axis represents log2-fold changes and Y-axis negative log10 p-values. Small values (1e-293) are added to p-values to avoid the underflow.

To verify that Kanade reconstructed meaningful counts, expressions of cell type markers and condition-dependent genes were visualized in Fig. 3D. In the reconstructed counts, the markers of each cell were expressed in the corresponding cells, and the condition-dependent genes exhibited variations specifically within the corresponding cells. This result indicated that biologically meaningful counts were reconstructed. Next, to demonstrate that Kanade selectively corrected the batch effects, we performed differentially expressed gene (DEG) analysis between conditions on a specific cell type and drew two volcano plots. Using original counts, there was little separation between ground-truth DEGs and non-DEGs in the volcano plot but both were separated using reconstructed counts (Figs. 3E and F). Moreover, using original counts, in spite of small log-fold changes, smaller p-values than true DEGs were detected when cell-specific marker genes (Green triangles in Fig. 3E) were affected by batch effects. In contrast, using reconstructed counts, this effect was mitigated, and marker genes showed higher p-values than true DEGs (Fig. 3F). These results indicated that Kanade distinguished batch effects from condition-dependent variations and corrected them selectively.

When Kanade was applied to Continuous data, mean reconstructed counts of genes in each cell type and time point were correlated to ground truth in the simulation (Fig. S2A). When dimensionality reduction on reconstructed counts was performed, cells were ordered along the time-series whereas batches in each time point were mixed (Figs. S2B and C). Both LISIs for the batch IDs and for time points in reconstructed counts (PCA Reconst in Figs. S1H and I) showed similar distributions to those in counts without batch effects (PCA woBE in Figs. S1H and S1I). Moreover, in reconstructed counts, cell-type-specific marker genes were expressed specifically in corresponding cell types, and time-dependently-regulated genes showed gradual variations over time (Fig. S2D). In addition, as well as Discrete data, the effect in which DEG analysis calculated small p-values of cell-type-specific marker genes by using uncorrected counts was suppressed when using reconstructed counts (Fig. S2E and F). These results demonstrated that reconstruction by Kanade was performed well also on the Continuous data.

### Application of Kanade for diabetes datasets

To verify the performance of Kanade with real data, we applied Kanade to the scRNA-seq atlas of the Mouse pancreas (Fig. 4A) [14–19]. This atlas consists of multiple datasets. They include pancreatic cells from control mice, type I (NOD) and type II (db/db) diabetes model mice, and chemical-stressed mice. Cells from db/db model mice with some treatment, vertical sleeve gastrectomy (VSG), and pair-fed (PF) were also included. The ages of mice in this atlas are not uniform. Because the ages of mice in this atlas were highly confounded with the original dataset, a major sources of the batch effect, when we applied Kanade to the atlas, a latent variable to predict the age of mice was prepared in addition to a set of latent variables to predict conditions. This variable was averaged when batch correction was performed. To explore the raw data structure, we generated the UMAP plot based on uncorrected counts and found some clusters were separated depending on batches or original datasets in spite of the same conditions and cell types (Fig. 4B and C). As the input cell types, we roughly cluster cells using Harmony, ignoring conditions (Fig. S3A and B, see Methods).

**Fig. 4:**
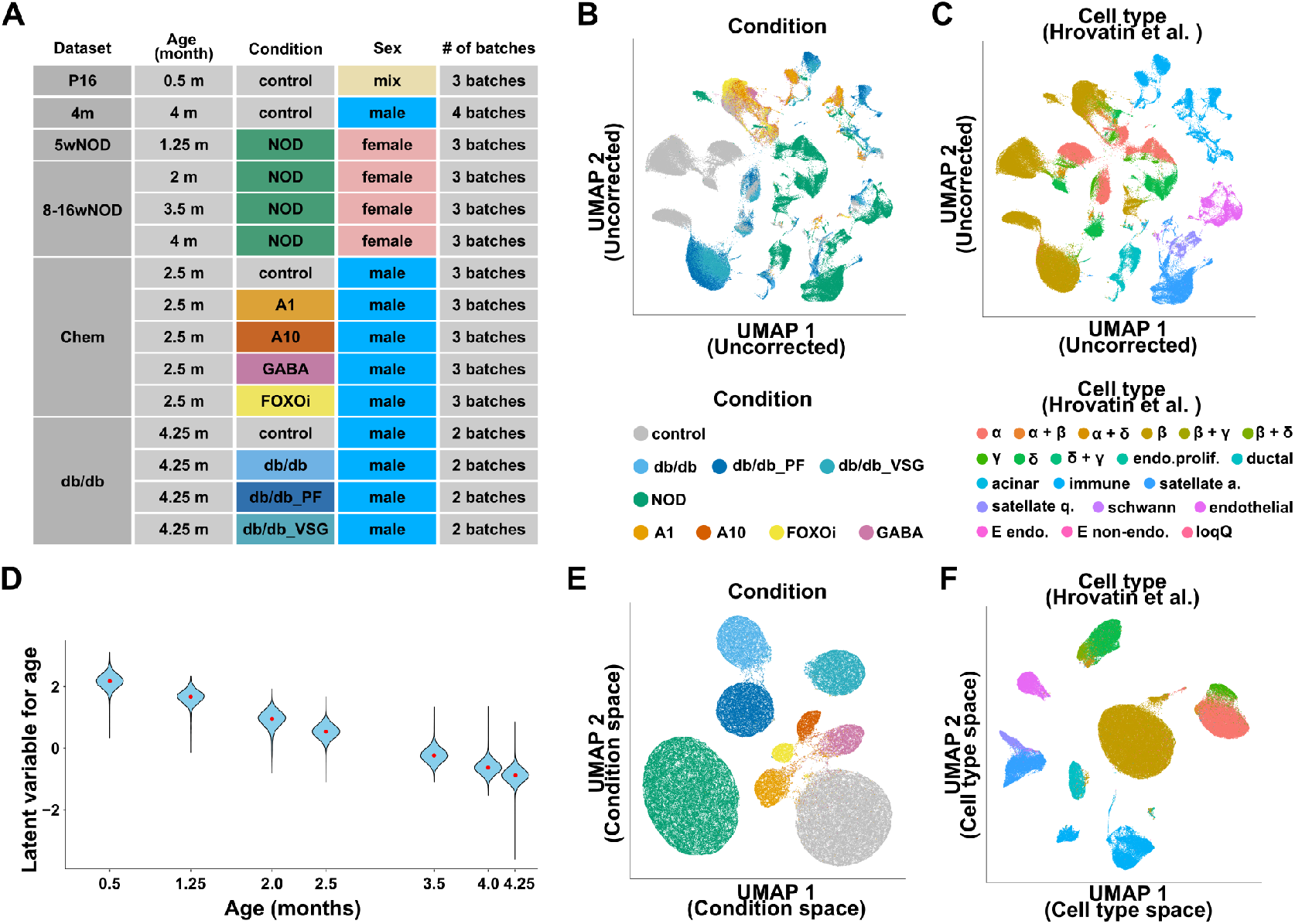
Demensionality reduction of mouse pancreatic atlas. (A) Overview of the pancreatic atlas used in this study from Hrovatin et al. (B and C) UMAP plots of the pancreatic atlas based on log-normalized counts. The colors of dots show the condition (B) and cell type annotated by Hrovatin et al. (C) of the cells. (D) Violin plot showing the distribution of the latent variable for age at each time point. The x-axis represents age in months, with each violin positioned accordingly. Red dots indicate the median values at each time point. (E) UMAP plot of the pancreatic atlas based on latent variables for conditions. The colors of the dots show the conditionthe of cells. Colors are the same as (B) (F) UMAP plot of the pancreatic atlas based on latent variables for cell types. The colors of the dots show the cell type annotated by Hrovatin et al. of cells. Colors are the same as (C) A1 and A10: artemether (1 or 10 μM); FOXOi: FoxO inhibitor endo.: endocrine; prolif.: proliferating; stellate a.: stellate-activated; stellate q.: stellate-quiescent; low-Q: low quality cell; E: embryonic (misannotated) *α* + *β, α* + *δ, β* + *γ, β* + *δ* and *δ* + *γ* means doublet of two cell types.

First, we found the encoder of Kanade was able to separate the signals in the latent variable space. The medians of latent variables for ages were clearly correlated with ages of input labels whereas batches from the same time points showed small differences among them (Figs. 4D and S4). In the latent variable space for conditions, cells were separated depending on the conditions, and batches in the same conditions were mixed (Figs. 4E and S5). Importantly, control cells from different datasets were mapped into the same clusters (Fig. S5A). In the latent variable space for cell types, cells were separated by the input labels but not batches (Figs. S3C and S6). Interestingly, we found more detailed cell types than input labels were distinguishable in the space (Figs. 4F and S7A). For example, alpha cells and gamma cells were separated in the UMAP plot in spite of both being labeled as input cluster 1. Moreover, lymphoid cells (input cluster 3) were separated into B cells, plasma cells, T cells, and the cell-cycling population of each cell type in the UMAP plot (Fig. S7A). These results suggest that Kanade is robust for a coarse input about cell types and ine-grained annotations are not necessary.

Next, we validated whether reconstructed counts contained biological features. In the UMAP plot based on the reconstructed counts, cells were clustered based on cell types and conditions, and no batch-specific clustering was observed (Figs. 5A-C, and S8). Note that control cells were mapped into the same clusters regardless of the source dataset as well as the latent variable space (Figs. S8). This suggests that batch effects were corrected well. Each cell type expressed the corresponding marker gene, suggesting the reconstructed counts retained distinctive cell type features (Fig. 5D). It is important that, as in the case of the latent variable space, more detailed cell types than input labels were distinguishable in the reconstructed counts. For example, potential doublet cells of pancreatic alpha cells and beta cells co-expressed glucagon (*Gcg*) and insulin I (*Ins1*). In addition, lymphoid cells (B cells, T cells, and plasma cells) showed cell-type-specific gene expression patterns [20–26].

**Fig. 5:**
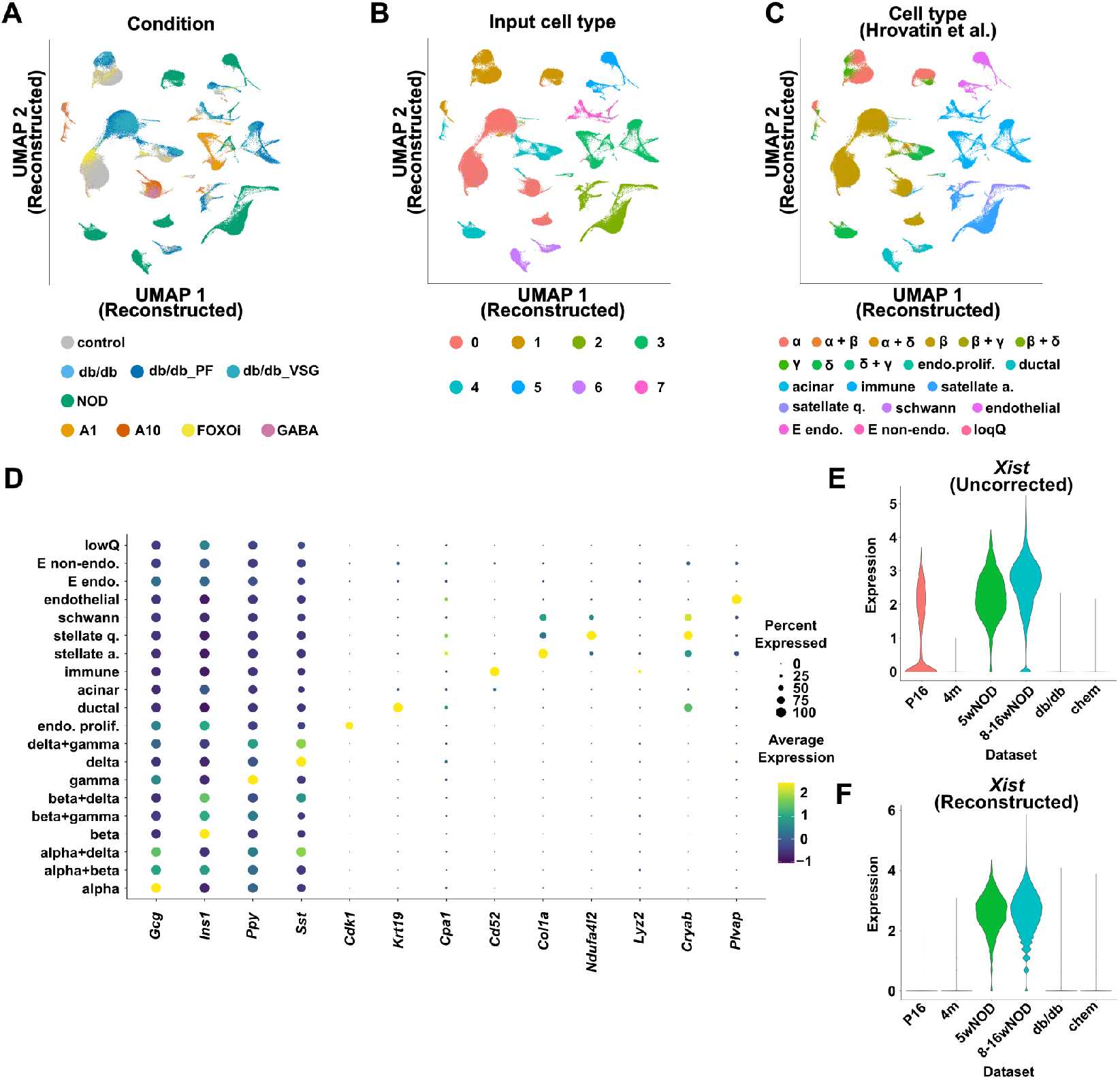
Reconstructed counts of the mouse pancreatic atlas. (A-C) UMAP plots of pancreatic atlas based on reconstructed counts. The color of the dots show conditions (A), input cell types (B), and cell types annotated by Hrovatin et al. (C). (D) Dot plot of the expression marker gene of each cell type shown in Hrovatin et al. (E and F) Violin plots of expression of *Xist* in each dataset based on uncorrected counts (E) and reconstructed counts (F). A1 and A10: artemether (1 or 10 μM); FOXOi: FoxO inhibitor endo.: endocrine; prolif.: proliferating; stellate a.: stellate-activated; stellate q.: stellate-quiescent; low-Q: low quality cell; E: embryonic (misannotated) *α* + *β, α* + *δ, β* + *γ, β* + *δ* and *δ* + *γ* means doublet of two cell types.

To demonstrate whether Kanade was able to selectively correct batch effects and keep conditional features, we focused on the expression of Inactive X specific transcripts (*Xist*). This gene is the one expressed only in female [27]. In the atlas used in this study, most cells are of male origin, while all cells in the NOD condition and only some of the control cells from the P16 dataset are of female origin (Fig. 4A). Consistent with this, *Xist* was detected in all batches under the NOD condition and only in batches derived from P16 under the control condition, but hardly detected in other batches under the control condition or any batches under other conditions before the correction (Fig. 5E). In this context, *Xist* expression can be regarded as a pseudo-NOD-specific feature and its expression in P16 dataset cells arose from dataset-specific differences, allowing it to be considered a batch effect. Surprisingly, in the reconstructed counts, *Xist* expression in cells from the P16 dataset almost disappeared, while it was maintained in cells from the two datasets under NOD condition (Fig. 5F). This result indicated that Kanade was able to distinguish batch effects and condition-specific features and correct only batch effects selectively.

Furthermore, to demonstrate the ability of Kanade to conserve conditional features in a diabetic context, we focused on pancreatic beta cells. First, expressions of diabetes-associated marker genes shown in Hrovatin et al. were examined (Fig. 6A). In the NOD condition beta-2 microglobulin (*B2m*), a type I diabetic marker [28], showed higher expression than other conditions, and three db/db conditions showed higher expression of type II diabetic markers: aldehyde dehydrogenase family 1, subfamily A3 (*Aldh1a3*) [29], vitamin D binding protein (*Gc*) [30], cholecystokinin (*Cck*) [31], and Gastrin (*Gast*) [32]. Next, principal component analysis (PCA) was performed on reconstructed counts of beta cells and principal component (PC) 3 and PC 2 separated the NOD condition and the db/db conditions respectively (Fig. 6B). GO terms related to the immune system including “innate immune response” (GO:0045087) were enriched in the top 100 genes that negatively contributed to PC 3 (Fig. 6C). These results were consistent with the fact that type I diabetes is an autoimmune disease [33]. GO terms related to the response to nutrition and diet including “response to food” (GO:0032094) were enriched in the top 100 genes that negatively contributed to PC 2 (Fig. 6D). In addition, PC 2 and PC 6 separated db/db VSG from db/db mice, but no PC was found which separated db/db PF conditions (Fig. 6E). In fact, expressions of *Cck* and *Gast*, genes with up-regulation in db/db conditions, were significantly downregulated in db/db VSG compared to db/db (Fig. 6F and 6G). These are consistent with the original paper of the dataset, which showed VSG results in a beta cell recovery from the diabetic characters but not PF [18]. Therefore, Kanade was able to conserve the features in the diabetic context.

**Fig. 6:**
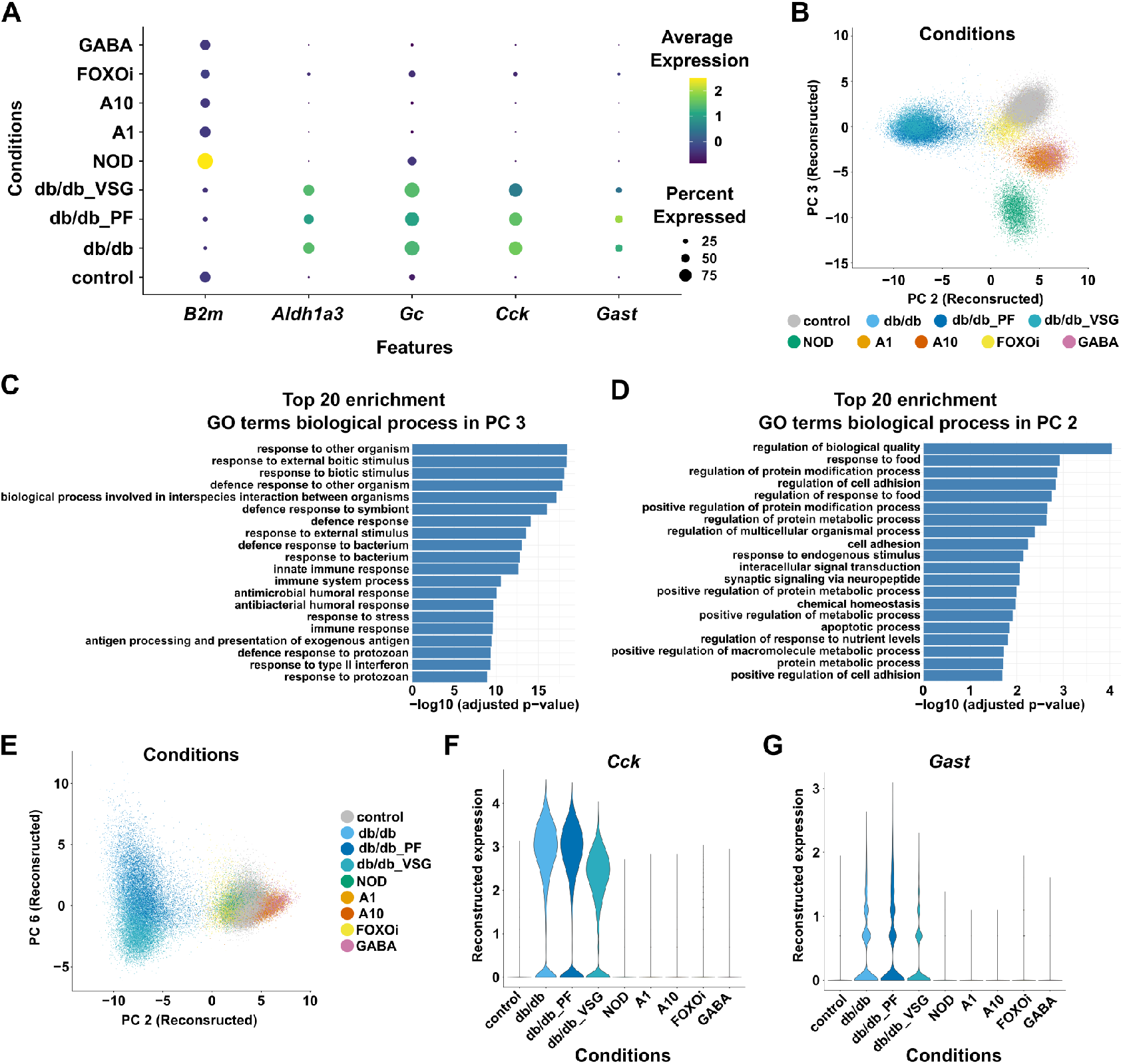
Conservation of conditional effects among conditions in the reconstructed counts of beta cells. (A) Dot plot of expressions of marker genes of type I diabetes and type II diabetes in beta cells. (B) PCA plot of beta cells. The X-axis and the Y-axis represent PC 2 and PC3 respectively. The colors of the dots show the conditions of cells. (C and D) Results of GO term enrichment analysis for Top 100 genes negatively contributed to PC 3 (C) and PC 2 (D). Barplots show negative log values of adjusted p-values. (E) PCA plot of beta cells. The X-axis and the Y-axis represent PC 2 and PC6 respectively. The colors of the dots show the conditions of cells. (F and G) Violin plots of expression of Cck (F) and Gast (G) in each dataset based on reconstructed counts. A1 and A10: artemether (1 or 10 μM); FOXOi: FoxO inhibitor

## Discussion

We developed Kanade, a VAE-based method to disentangle condition-specific features from batch effects and to selectively correct the latter. For both simulated and real datasets, we demonstrated that Kanade separated condition-specific features and batch effects in latent variable spaces and preserved condition-specific features in the reconstructed and corrected counts. Comparisons between conditions are fundamental to the biological research and a correction method dedicated to this purpose has been an important need. Our method is expected to meet that need and be applied to research in medicine and drug discovery, and more fundamental regions including developmental biology.

A key characteristic of Kanade is its ability to explicitly separate information by assigning predictors of specific attributes to designated subspaces in the latent variable space. This approach improves both interpretability and practical applicability. It is expected that Kanade can be extended by integrating it with various statistical or machine-learning models, further enhancing its applicability and performance. Moreover, unlike existing batch correction methods such as Harmony, which primarily focus on embedding data into a shared latent space, Kanade leverages the advantages of generative modeling to achieve batch correction in the reconstructed expression space. This feature enhances interpretability, thereby improving accessibility for biological researchers. In addition, the generative nature of Kanade, combined with its ability to assign specific latent variables to designated attributes, enables potential applications such as interpolating the state between conditions.

However, Kanade has certain limitations. Notably, it is unable to handle previously unseen batches and cannot disentangle dataset-wide shared information, such as variations arising from differences between laboratories. It is possible that training on a larger and more diverse dataset may allow the model to evolve into a more robust foundation model capable of handling broader datasets. In summary, Kanade offers a flexible approach to selectively correcting batch effects while preserving condition-specific information and balancing interpretability and usability in batch correction in single-cell omics.

## Acknowledgments

This research was supported by the Moonshot R&D–MILLENNIA Program (grant number JPMJMS2024-9; to H.N.) and the Cooperative Study Program of Exploratory Research Center on Life and Living Systems (ExCELLS; program number 19–102 to H.N.). We would like to thank Susei Fujioka for valuable advice on marker genes of immune cells, Ryosuke Kojima for advice on the model development, and the members of Honda Lab and other laboratories of the mathematical biology group at Hiroshima University for their insightful discussion. Additionally, we acknowledge the assistance of ChatGPT in coding and manuscript writing.

## Author contributions

S.S., M.T., K.N. and H.N. conceived the model. S.S. and M.T. developed the software. S.S. performed validations and applied the model to real data. S.S., M.T., K.N, and H.N. wrote the manuscript. H.N. supervised the work.

## Methods

### Kanade model

Kanade assumed *x*_*g,i*_, the gene expression count of gene *g* in cell *i*, is sampled from the zero-inflated negative binomial (ZINB) distribution defined by latent variables. This process can be written down as follows:

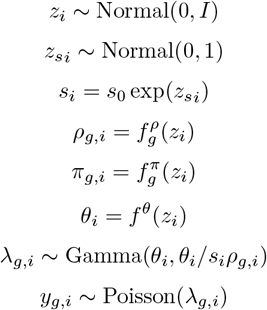

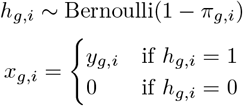

The prior distributions of latent variables *z*_*i*_ ∈ ℤ^*K*^ and 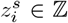 are assumed to be independent normal distributions. The scale factor *s*_*i*_ is calculated as an exponential of 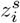. From *z*_*i*_ the parameters of ZINB *ρ*_*g,i*_ and *θ*_*g,i*_ are calculated by function 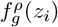 and 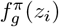 respectively. These functions are modeled by a neural network (NN decoder) and outputs are scaled by softmax function for *ρ*_*g,i*_ and logistic function for *π*_*g,i*_ respectively. The parameter *θ*_*i*_, common in all genes, is calculated by a function 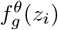, the softplus function of the weighted sum of elements of *z*_*i*_.

The parameter *λ*_*g,i*_ of Poisson distribution is sampled from the Gamma distribution with shape parameter *θ* and rate parameter 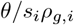, and a true count *y*_*g,i*_ is sampled from Poisson distribution with the sampled *λ*_*g,i*_. To inflate zero-values, the count *x*_*g,i*_ is forced to zero with probability *π*_*g,i*_, otherwise *x*_*g,i*_ = *y*_*g,i*_.

Kanade modeled this process as a variational autoencoder-based model using PyTorch [34]. The input of Kanade is *x*_*i*_ and the encoder neural network, NN encoder, projected the input into means *mu* and variances *σ*^2^ (*σ*^2^ is log-scaled) of latent variable *z*_*i*_ and *z*_*si*_. *z*_*i*_ and *z*_*si*_ are sampled using the parameterization trick based on calculated mean and variance. From *z*_*i*_, ZINB parameters are calculated in the way explained above. In this paper, the NN encoder had three layers and the numbers of nodes were 2000, 400, and 200 from input respectively. Each layer consisted of batch normalization, ReLU activation, and dropout sublayers in this order. NN decoder had three layers with the same sublayers as NN encoder, and the numbers of nodes were 200, 400, and 2000 from input respectively.

In Kanade model, a predictor Pred *l* is provided to predict the label *l* to be considered. The Pred *l* is a linear predictor that predicts the label *l* from the set of elements 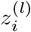 of *z*_*i*_. Each 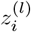 has no common elements with each other. For the continuous label (time point of Continuous data and the ages of mice in the pancreatic atlas), the predictor functions as a regressor by computing a weighted sum of 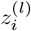, whereas for discrete labels (the other labels), it acts as a classifier by applying a softmax function to the weighted sum of 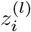.

### Model training scheme

The VAE part and predictors of Kanade were simultaneously trained using the loss function below:

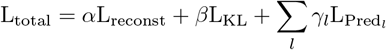

Here, L_reconst_is a reconstruction loss and is calculated as a negative log-likelihood.L_KL_is a KullbackLeibler divergence between prior and posterior probability of 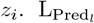 is a prediction loss (cross entropy loss for discrete labels and mean squared error for continuous labels) of Pred *l. α, β* and *γ*_*l*_ are weights of each loss. The used weights in this study are listed in Table.S1.

### Training for simulated datasets

The numbers of assigned elements of latent variables were one for the conditions, three for cell types, and five for batches. In our training, the minibatch size is 128 and the number of epochs is 25. Adam optimizer was used for all training and the learning rate is 0.001. The dataset was split into training and test data at a 3:1 ratio, and parameters and seeds were selected for which the model had small overlearning.

### Training for mouse pancreatic atlas

The numbers of assigned elements of latent variables were one for the age of mice, three for conditions, five for cell types, and 10 for batches. In our training, the minibatch size is 128 and the number of epochs is 50. Adam optimizer was used for all training and the learning rate is 0.001. The dataset was split into training and test data at a 3:1 ratio, and parameters and seeds were selected for which the model had small overlearning.

### Generation of simulated datasets

To evaluate Kanade, two simulated datasets were generated. The first dataset had discrete conditions (Discrete data, Fig. 2A). This dataset simulated an experiment comparing healthy wild-type and disease mutant. In the “Disease” condition, condition-dependent genes were up- or down-regulated compared with “WT”. This dataset contains two conditions, each with three batches, resulting in a total of six batches. The second dataset assumed that the expression of condition-dependent genes changes continuously over time and cells were sampled at five time points (Continuous data, Fig. S1A). Here, three batches were sampled at each time point and 15 batches were obtained in total. Both datasets include three cell types (A, B, and C). Batch effects were randomly introduced in each batch(Figs. 2A-C and S1A-C).

To reproduce this setting, we assumed a total of 20,000 genes and categorized them as follows: marker genes for each cell type (mA, mB, and mC); genes upregulated in each cell type under disease conditions (or over time) (uA, uB, and uC); genes downregulated in each cell type under disease conditions (or over time) (dA, dB, and dC) and other non-variable genes (others). For simplicity, these categories have no overlap. The numbers of each gene type are listed in Table S2. Variable genes were randomly chosen from 15,000 genes with the largest basic mean-expressions, which are explained next.

To generate simulated datasets, the method of Splatter [35] was modified. First, basic mean-expressions of 20,000 genes were sampled from a gamma distribution (shape = 0.6 and rate = 0.3). For outlier expression, genes were sampled from 20,000 genes using a binomial distribution with a mean 0.05, and the basic mean-expression of each sampled gene *g* was replaced by *m*_*g,out*_ defined below,

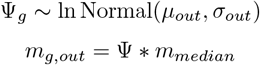

Here, *μ*_*out*_ = 4 and *σ*_*out*_ = 0.5 were used, and *m*_*median*_ was the median value of basic mean-expression of all genes before the replacement.

To introduce cell-type-dependent variation, the magnitude of variation for a variable gene *g* was sampled by

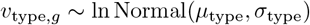

Here, *μ*_type_ = 2 and *σ*_type_ = 0.5. If *v*_type,*g*_ *<* 1.1, the value was replaced by 1.1 to ensure a minimum variation threshold.

- For marker genes and downregulated genes, the mean expression of *g* was *v*_type,*g*_ folded in the corresponding cell type and 1*/v*_type,*g*_ folded in the other cell types.
- For upregulated genes, the mean expression was 1*/v*_type,*g*_ folded in the corresponding cell type.

To account for condition-dependent variation, magnitudes of variation for a condition-dependent gene *g* was sampled by

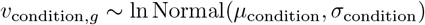

Here, *μ*_condition_ = 2 and *σ*_condition_ = 0.5. If *v*_condition,*g*_ *<* 1.1, the value was replaced by 1.1 to ensure a minimum variation threshold.

- For Discrete data:

- If *g* was the upregulated gene, its mean expression was *v*_condition,*g*_ folded in the corresponding cell type in the disease condition.
- If *g* was the downregulated gene, its mean expression was 1*/v*_condition,*g*_ folded.
- For Continuous data:
- If *g* was the upregulated gene, its mean expression of upregulated at time point *t* was 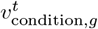 folded
- If *g* was the upregulated gene, its mean expression of upregulated at time point *t* was 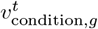 folded, and if the downregulated gene 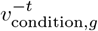 folded.

For a representation of batch effects, Random perturbations were applied to the mean expression levels of genes for each batch. Specifically, for batch *b* and gene *g, c*_*g,b*_ was randomly assigned such that it took the value 0 with a probability of 0.8, 1 with a probability of 0.1, and -1 with a probability of 0.1. Additionally, the magnitude of the batch effect of gene *g* was sampled from a log-normal distribution:

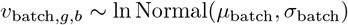

Here, *μ*_batch_ = ln 2, *σ*_batch_ = 0.12. If *v*_batch,*g,b*_ was less than 1.1, it was replaced with 1.1 to ensure a minimum variation threshold. Using these values, the mean expression level of gene *g* was 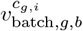 folded. For the simulated dataset without batch effects, this step was skipped.

To capture single-cell variability, the following procedure was implemented. Let 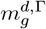 denote the mean expression level of gene *g* under condition d in cell type Γ(∈ *{A, B, C}*), as defined above. First, the library size ŝ_*i*_ for each cell *i* was sampled using the following equation:

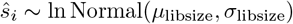

Here, *μ*_libsize_ = ln 5000 and *σ*_libsize_ = 0.2. Using these values, the expression level 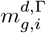, adjusted for the library size of each cell, was computed as follows:

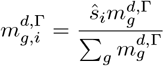

True expression value of gene *g* in each cell 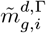 was obtained by following:

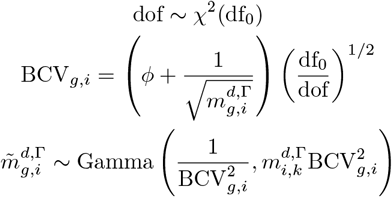

Here, df_0_ = 60 and *ϕ* = 0.1. This process was performed for each batch and cell type.

To simulate RNA-seq sampling, Poisson distribution was used:

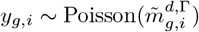

Here, *y*_*g,i*_ was the true count of a gene *g* in a cell *i*. Usually, small library sizes of single-cell RNA-seq result in dropout, where some gene count values are artificially recorded as zero despite the gene being expressed. Thus, the observed value *x*_*g,i*_ was obtained as follows:

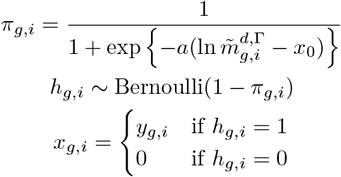

The parameters were set to *a* = −1 and *x*_0_ = 0.

### Preprocessing of mouse pancreatic islet scRNA-seq atlas data

The mouse pancreatic islet scRNA-seq atlas [14] was downloaded from CELLxGENE database (id: 296237e2-393d-4e31-b590-b03f74ac5070) as a h5ad file. From the atlas, three datasets were excluded for the following reasons: “mSTZ” dataset had only one batch per condition, most of the cells in “Embryonic” dataset were annotated as “unknown”, and “Aged” dataset had a large gap of age from other datasets. As the units of ages of samples varied in the original annotations, all were standardized to months. As some annotations were made with wide ranges, they were assigned the midpoint values. Condition labels were added depending on a combination of pathogenic stresses (genotypes or chemicals) and treatments (Table S3).

Annotation of cell-type labels was performed by Seurat v4 [8]. The original counts were loaded into a Seurat object, and the object was separated by the “donor id” labels, which corresponded to batch identifiers. Separated objects were normalized by SCTransform [36] function independently and merged again. The merged dataset was projected into the low dimensional space by RunPCA function and 30 principal components (PCs) were used for RunUMAP function. PCs were corrected by Harmony [9]. LISIs (see below) were calculated using 10, 30, and 50 Harmony-corrected dimensions, and we decided to use 10 dimensions for the clustering and UMAP projection because the majority of cells had LISI values larger than one. Clustering was performed by FindNeighbors and FindClusters function (“resolution” parameter was set to 0.1) functions and its result was used for Kanade training as cell-type labels. Marker genes of each cluster were detected by FindAllMarkers function and each cluster was confirmed to correspond to a specific cell type. After annotations, genes which expressed only in less than ten cells were excluded for the stable learning of models.

For the training, original counts were used. Before training, genes expressed in less than 10 cells were removed.

### Count reconstruction and downstream dimensionality reduction

For simulated data, uncorrected counts were input to the trained encoder and projected to latent variables. After latent variables for the batches were replaced with zeros, ZINB parameters were obtained from latent variables by the trained decoder. Among these parameters, the scale factors were averaged, and counts were reconstructed based on the obtained ZINB parameters. For the pancreatic atlas, in addition to the replacement of latent variables for the batches, latent variables for the ages were averaged before decoding. Downstream analysis was performed by Seurat v4. The reconstructed counts were log-transformed after the addition of 1. Using FindVariableFeatures function, 3,000 highly variable genes (HVGs) were detected. After reconstructed counts of HVGs were standardized with ScaleData function, PCA was performed with RunPCA functions. UMAP was performed with RunUMAP function and the top 30 PCs. For the analysis of beta cells in the pancreatic atlas, Cells with the label “beta” of the metadata “cell type reannotatedIntegrated” by Hrovatin et al. after the analysis above. PCA was re-performed on the extracted beta cells.

### Differentially expressed gene analysis

Differentially expressed gene analysis (DEG analysis) was performed by FindMarkers function in Seurat v4.. The parameter test.use was set to “MAST” [37]. For the simulated datasets, thresholding parameters “min.pct” and “logfc.threshold” were set to 0 so that all genes were considered. For the mouse pancreatic atlas, “logfc.threshold” was set to 0 and “min.pct” default value (0.01).

### Gene ontology (GO) term enrichment analysis

In the PCA of beta cells in the pancreatic atlas. The 100 genes with the smallest “feature.loading” values for PC 2 and PC 3 were listed respectively and used for the gene ontology (GO) term enrichment analysis. For GO term enrichment analysis, g:Profiler (https://biit.cs.ut.ee/gprofiler) [38] was used with g:SCS algorithm, and significantly enriched terms in cellular components were identified at an FDR threshold of 0.05 and term size lower than 5,000.

### Calculation of local inverse Simpson’s indices (LISI)

Preprocessing was performed by Seurat v4. and Harmony. Simulated datasets without batch effects and simulated uncorrected datasets with batch effects were normalized and scaled by NormalizeData function and ScaleData function with 3,000 HVGs detected by FindVariableFeatures function. Harmony applied to the top 30 PCs of uncorrected datasets. The preprocessing of reconstructed data was explained above. Here, the top 30 dimensions in PCs or Harmony-corrected variables were used for the calculation of local inverse Simpson’s indices (LISI). LISIs were calculated by the compute lisi function in R package lisi [9].

## Data and code availability

The code and data used for the analysis will be made publicly available after the paper is published. The mouse pancreatic islet scRNA-seq atlas by Hrovatin et al. is available from CELLxGENE database (id: 296237e2-393d-4e31-b590-b03f74ac5070). The Python package version of Kanade is scheduled to be released.

## Supplementary Figures

**Fig. S1:**
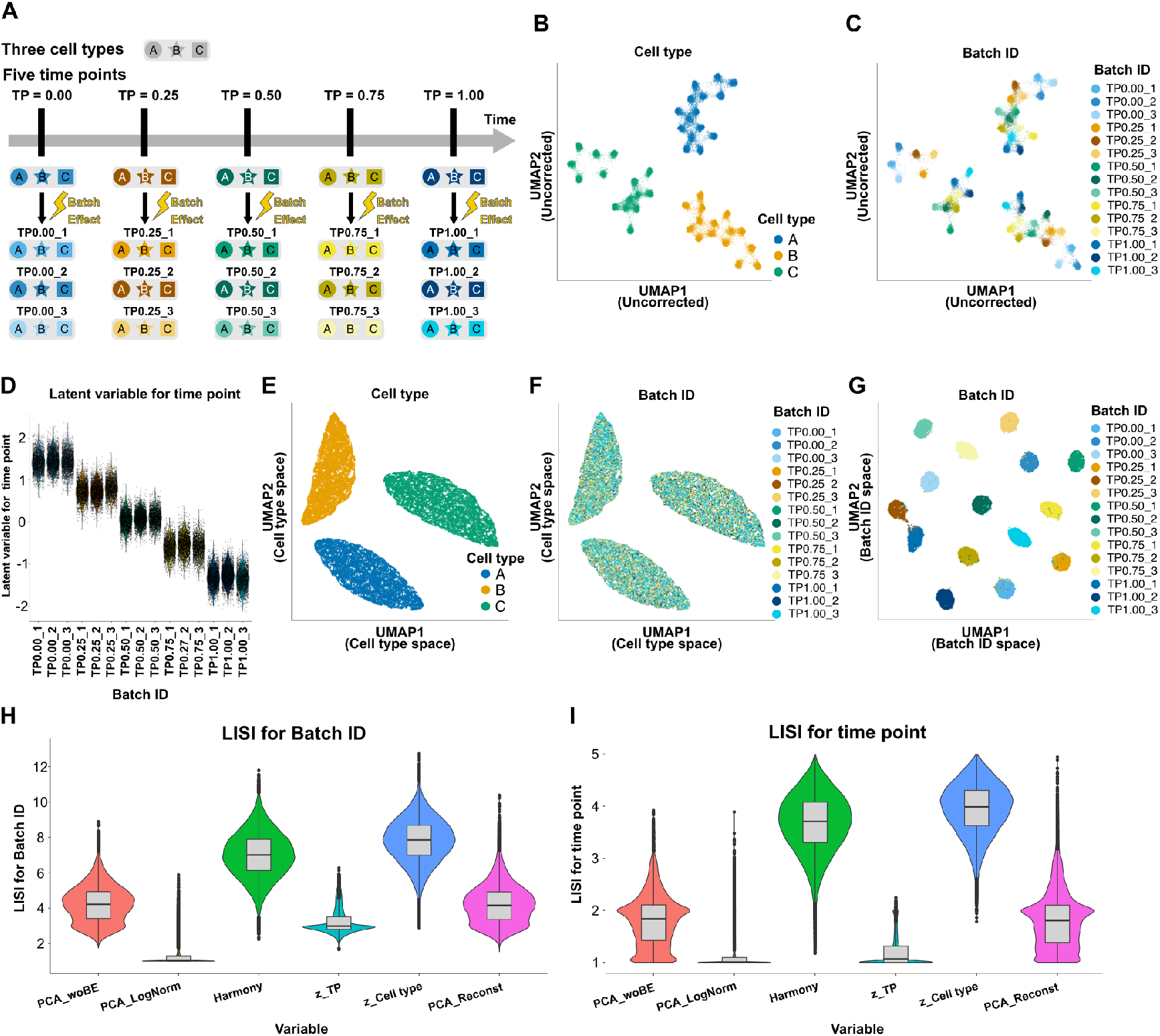
Simulated continuous data in the latent variable spaces of Kanade. (A) The scheme of Continuous data. (B and C) UMAP plots of discrete data based on log-normalization. The colors of the dots show cell types (B) and batch IDs (C). (D) The violin plots of the latent variable for time points across independent batches. Each dot represents an individual cell. (E and F) UMAP plots of continuous data using latent variables for cell types. The colors of the dots show cell types (E) and batch IDs (F).

**Fig. S2:**
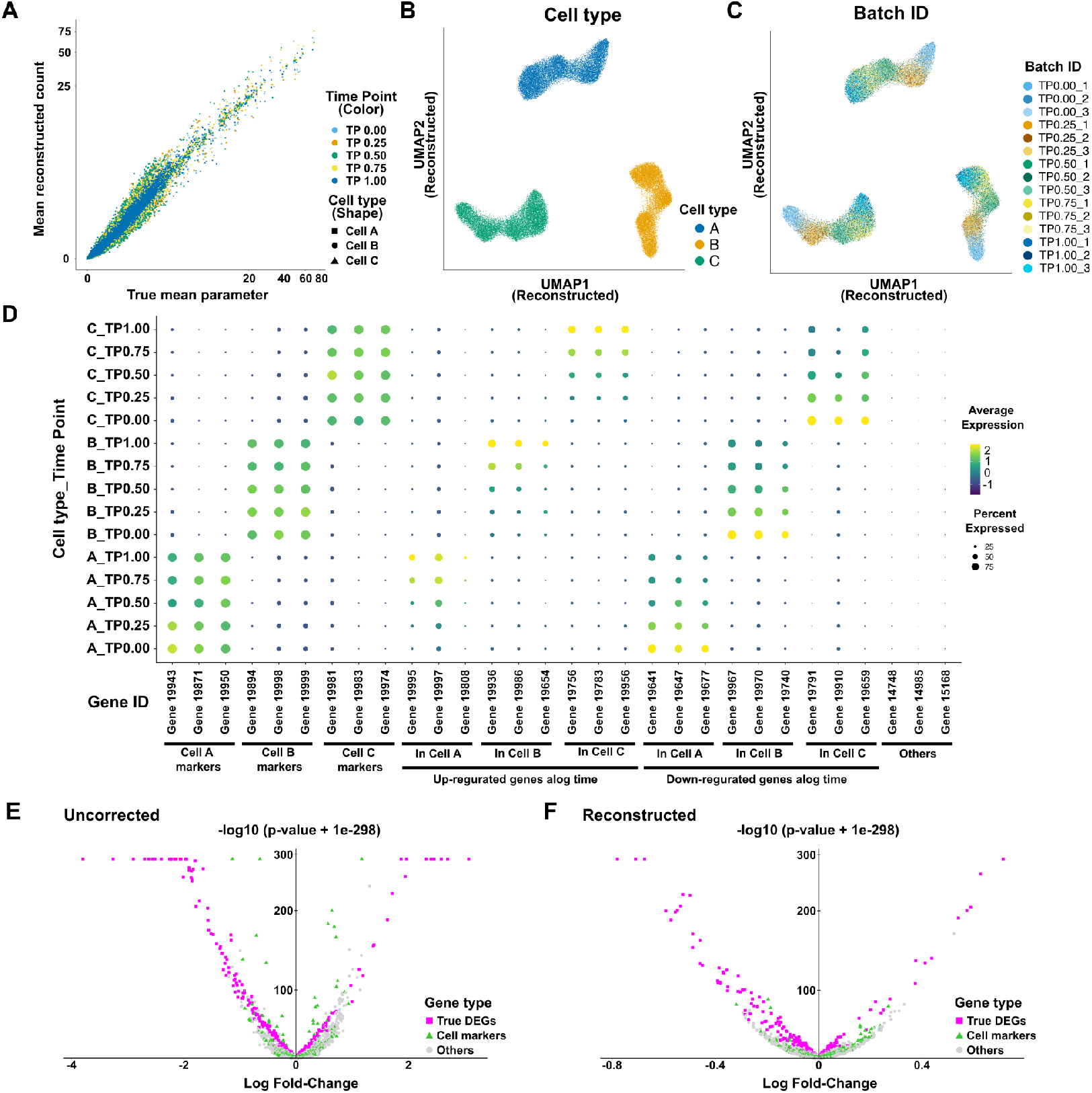
Reconstructed counts of simulated discrete data. (A) Scatter plots of mean gene expressions in each cell type and time point. The X-axis represents the mean parameters of the simulation and the Y-axis mean of reconstructed counts. Both axes are log-scaled. Each dot represents each gene, and the color and shape of the dots represent conditions and cell types respectively. (B and C) UMAP plots of continuous data based on reconstructed counts. The colors of the dots show cell types (B) and batch IDs (C). (D) The dot plot of reconstructed gene expressions in each cell type and condition. (E and F) Volcano plots of differentially expressed gene analysis results in cell type A between time points 0.0 and 1.0 based on uncorrected log-normalization (E) and reconstructed counts by Kanade (F). Each dot means a gene and X-axis represents log2-fold changes and Y-axis negative log10 p-values. Small values (1e-298) are added into p-values to avoid the underflow.

**Fig. S3:**
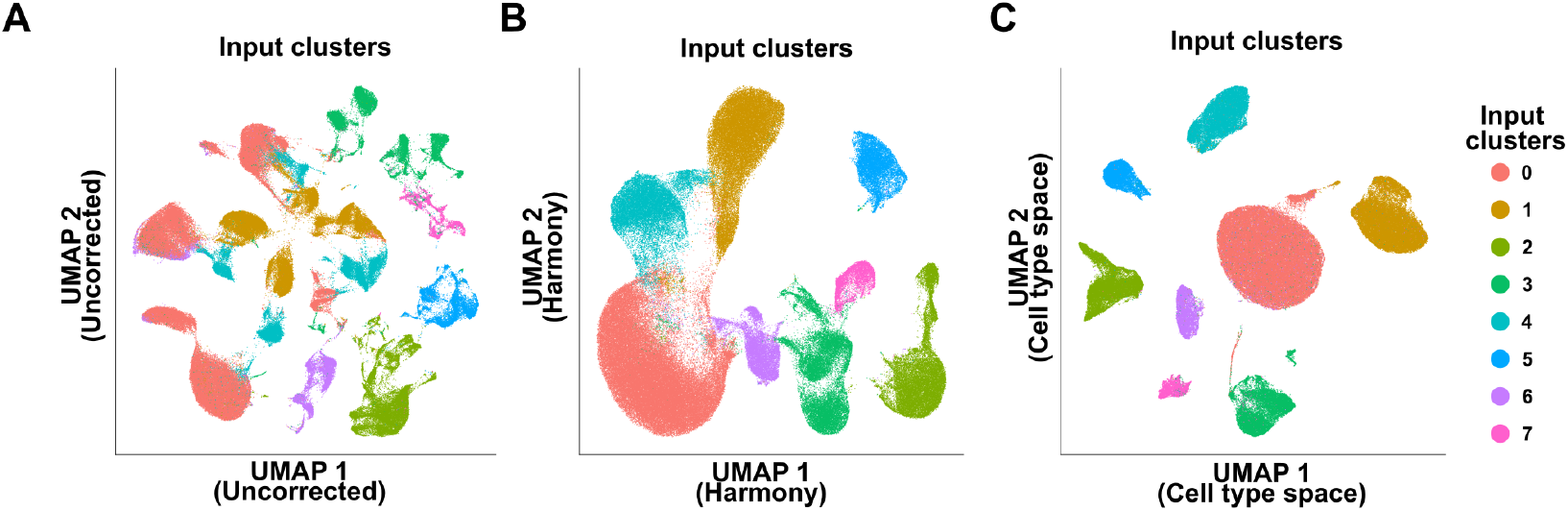
Input clusters of mouse pancreatic atlas for the training. Input clusters of mouse pancreatic atlas for the training of Kanade were shown with the color of dots on the UMAP plots based on log-normalization (A), Harmony-corrected (B), and latent variables for cell types of Kanade (C).

**Fig. S4:**
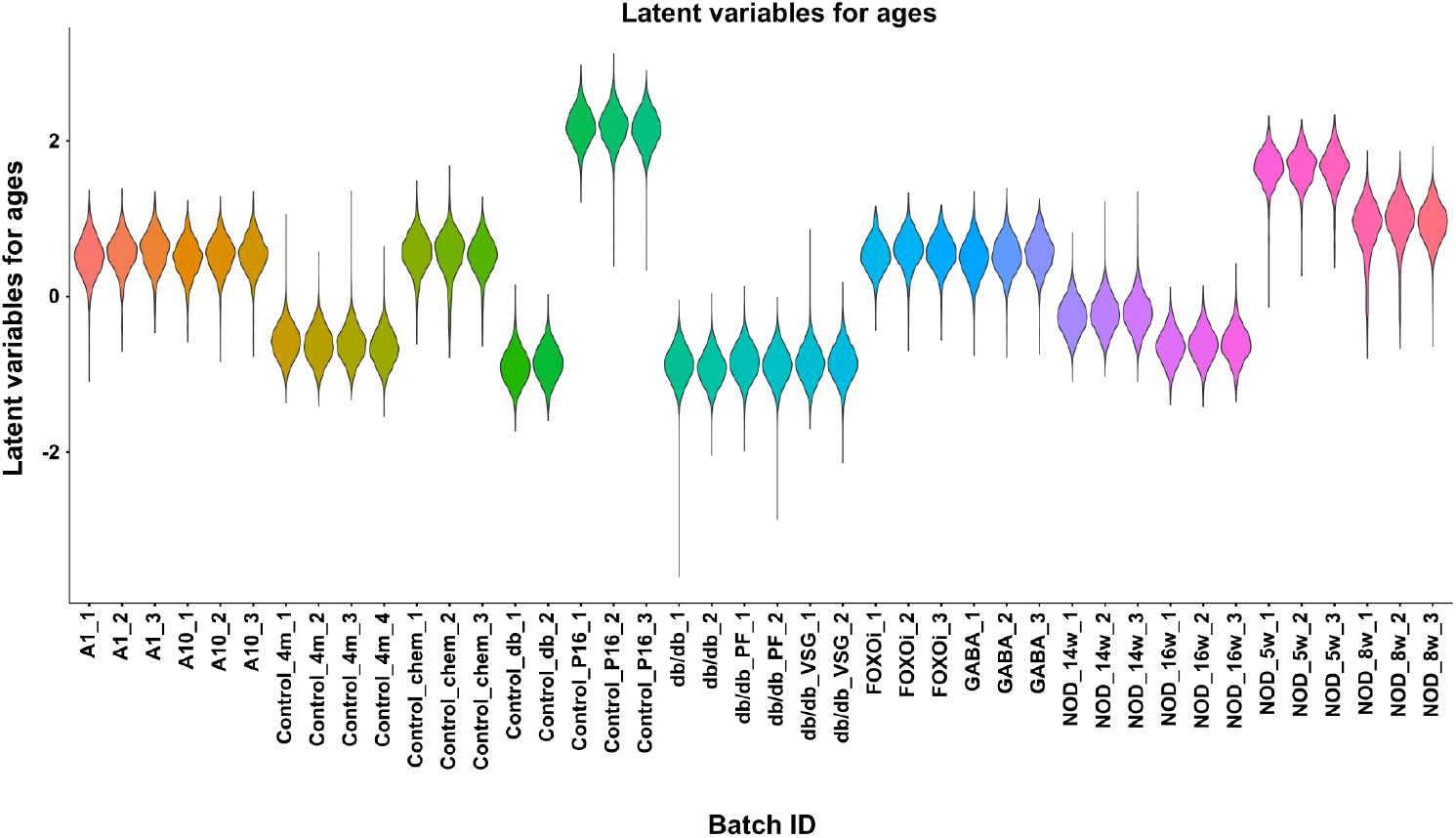
Latent variable for the ages of mice across the batches. Violin plots of the latent variable for the ages of mice across the batches.

**Fig. S5:**
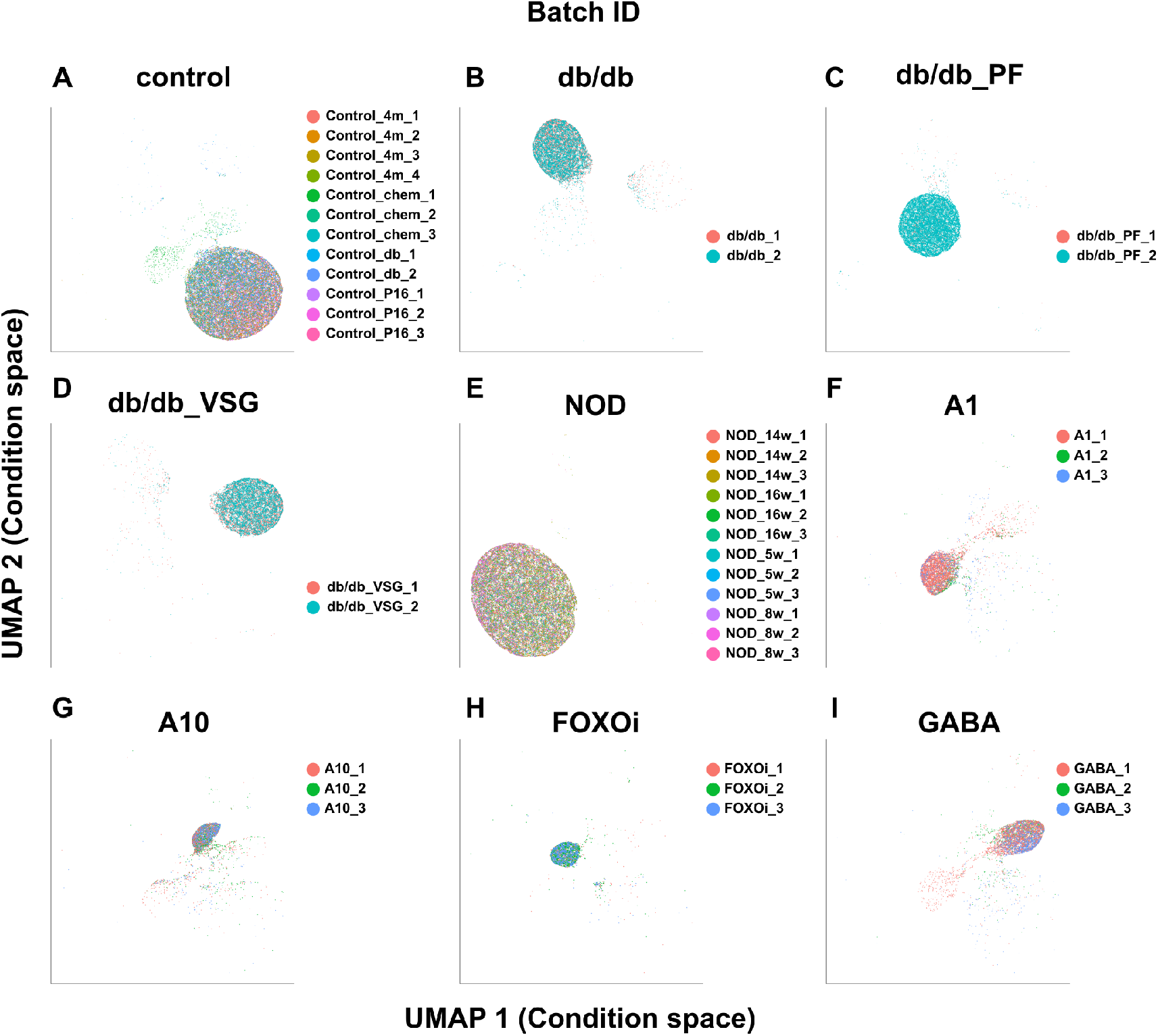
Batches on latent variables for conditions. UMAP plots of the mouse pancreatic atlas based on latent variables for conditions. The colors of the dots show batches, and plots were separated by conditions. (A) control, (B) db/db, (C) db/db PF, (D) db/db VSG, (E) NOD, (F) A1, (G) A10, (H) FOXOi and (I) GABA A1 and A10: artemether (1 or 10 μM); FOXOi: FoxO inhibitor

**Fig. S6:**
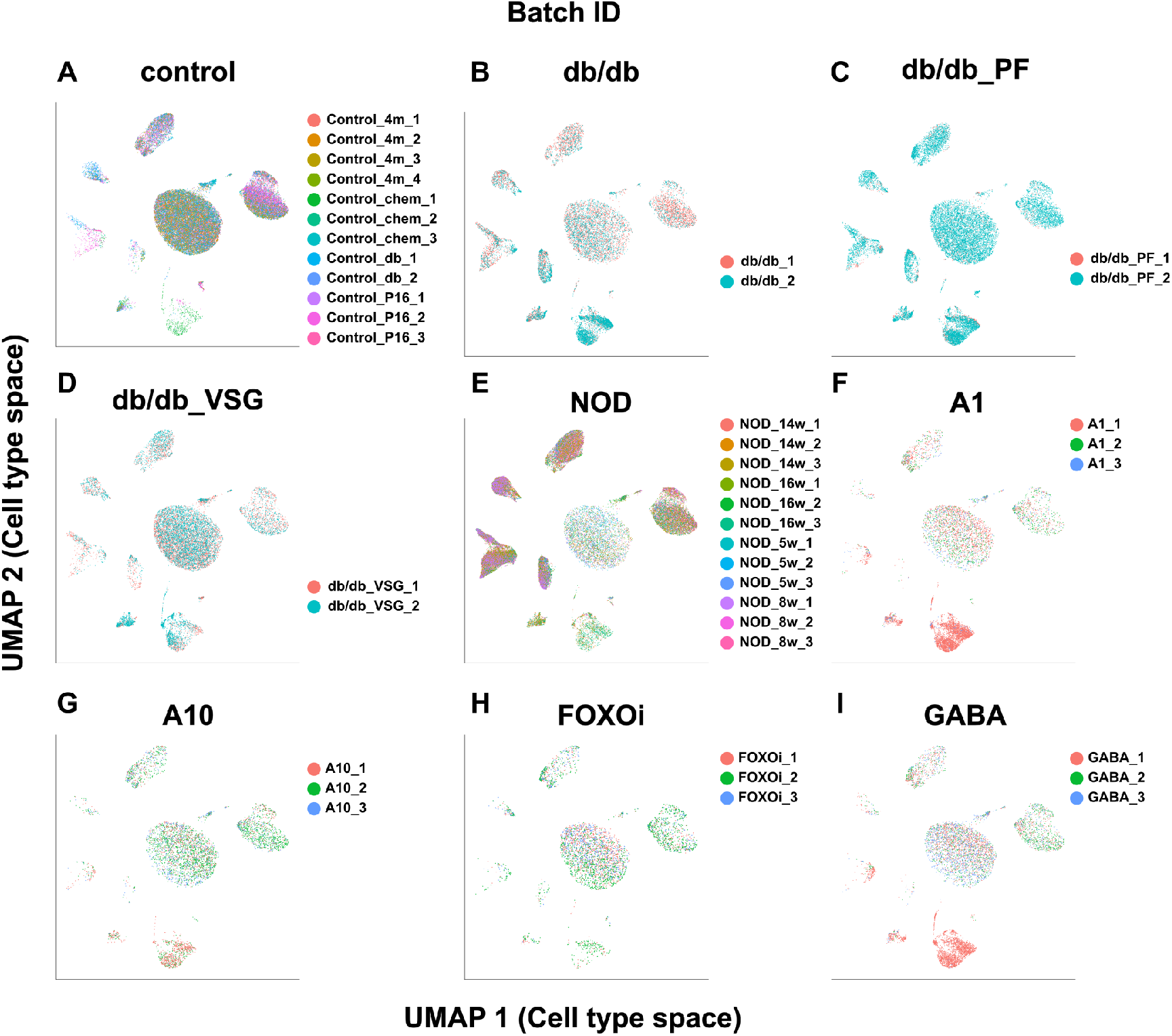
Batches on latent variables for cell types. UMAP plots of the mouse pancreatic atlas based on latent variables for cell types. The colors of the dots show batches, and plots were separated by conditions. (A) control, (B) db/db, (C) db/db PF, (D) db/db VSG, (E) NOD, (F) A1, (G) A10, (H) FOXOi and (I) GABA A1 and A10: artemether (1 or 10 μM); FOXOi: FoxO inhibitor

**Fig. S7:**
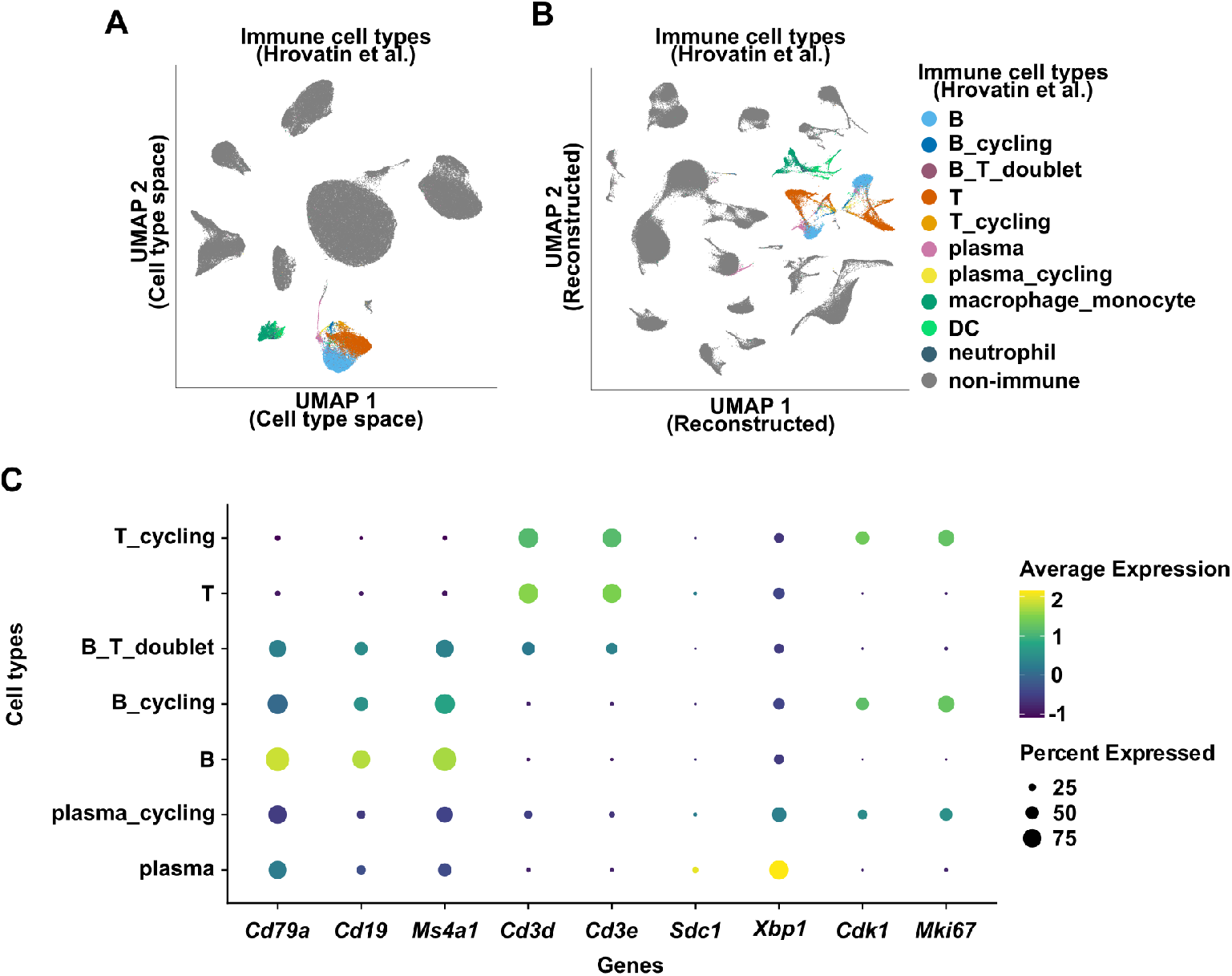
Separation of lymphoid cells. (A and B) UMAP plot of the mouse pancreatic atlas based on latent variables for cell types (A) and reconstructed counts (B). The colors of the dots show the annotation of immune cells by Hrovatin et al.. (C) Dot plots of expressions of lymphoid cell types in each cell type. Cell types were based on the annotation of immune cells by Hrovatin et al.. Only lymphoid cells were shown.

**Fig. S8:**
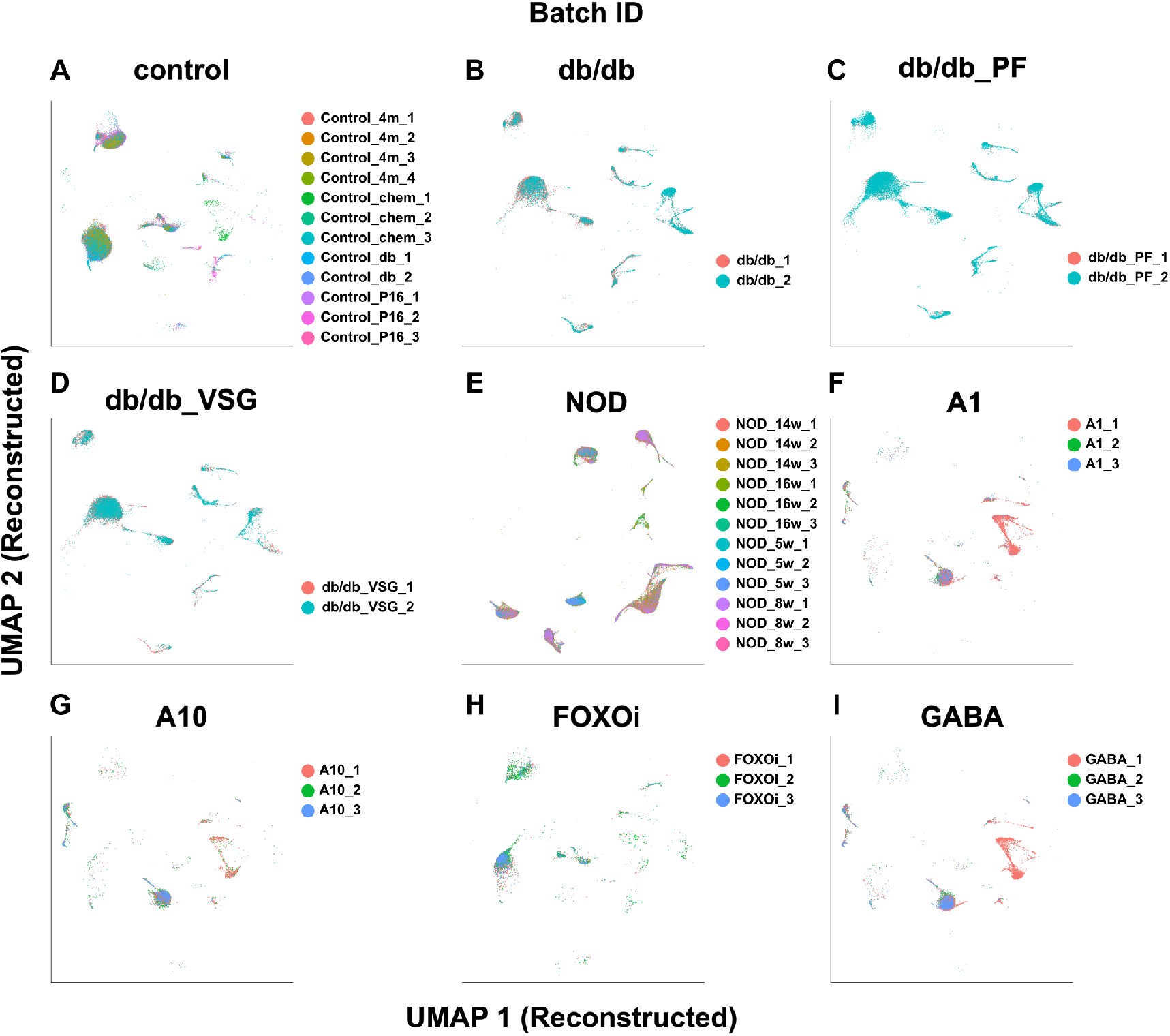
Batches on reconstructed counts. UMAP plots of the mouse pancreatic atlas based on reconstructed counts. The colors of the dots show batches, and plots were separated by conditions. (A) control, (B) db/db, (C) db/db PF, (D) db/db VSG, (E) NOD, (F) A1, (G) A10, (H) FOXOi and (I) GABA A1 and A10: artemether (1 or 10 μM); FOXOi: FoxO inhibitor

